# Evolution of a sex megachromosome

**DOI:** 10.1101/2020.07.02.182808

**Authors:** Matthew A. Conte, Frances E. Clark, Reade B. Roberts, Luohao Xu, Wenjing Tao, Qi Zhou, Deshou Wang, Thomas D. Kocher

## Abstract

Chromosome size and morphology vary within and among species, but little is known about either the proximate or ultimate causes of these differences. Cichlid fish species in the tribe Oreochromini share an unusual megachromosome that is ~3 times longer than any of the other chromosomes. This megachromosome functions as a sex chromosome in some of these species. We explore two hypotheses of how this sex megachromosome may have evolved. The first hypothesis proposes that it developed by the accumulation of repetitive elements as recombination was reduced around a dominant sex-determination locus, as suggested by traditional models of sex chromosome evolution. An alternative hypothesis is that the megachromosome originated via the fusion of an autosome with a highly-repetitive B chromosome, one of which had carried a sex-determination locus. Here we test these hypotheses using comparative analysis of several chromosome-scale cichlid and teleost genomes. We find the megachromosome consists of three distinct regions based on patterns of recombination, gene and transposable element content, and synteny to the ancestral autosome. A WZ sex-determination locus encompasses the last ~105Mbp of the 134Mbp megachromosome and the last 47Mbp of the megachromosome shares no obvious homology to any ancestral chromosome. Comparisons across 69 teleost genomes reveal the megachromosome contains unparalleled amounts of endogenous retroviral elements, immunoglobulin genes, and long non-coding RNAs. Although the origin of this megachromosome remains ambiguous, it has clearly been a focal point of extensive evolutionary genomic conflicts. This megachromosome represents an interesting system for studying sex chromosome evolution and genomic conflicts.

## Introduction

Almost two centuries of cytogenetic studies have revealed the great diversity of animal karyotypes. Chromosome numbers range from a single chromosome pair in the jack jumper ant (Crosland and Crozier 1986) to ~225 chromosome pairs in the Atlas blue butterfly (Lukhtanov 2015). Chromosome numbers can differ dramatically among closely related species even without changes in ploidy (Yang et al. 1997). While the total length of the chromosomes is directly related to genome size, and thus to factors such as population size (Lynch and Conery 2003), there is little theory to explain why chromosome numbers should vary so dramatically.

There are also dramatic differences among species in the shapes of chromosomes. Some lineages contain mostly acrocentric chromosomes, while others segregate mostly metacentric chromosomes. Differences among species have been attributed to supposedly random processes of chromosome fusion and fission. Recent data suggest that these differences may arise from changing biases in centromeric drive during female meiosis (Blackmon et al. 2019), but the molecular basis for these shifts remain obscure (Kursel and Malik 2018). We suspect there might be some common mechanisms and rules governing the variety of sizes and shapes of chromosomes in a particular lineage. However, at present we are unable to predict the structure of karyotypes – meaning the number and shape (acrocentric vs. metacentric) of chromosomes - in particular lineages.

The genome is a battleground on which genetic conflicts are fought on many levels. Selfish genetic elements such as transposons tend to proliferate, increasing the size of genomes (Canapa et al. 2016; Kapusta et al. 2017). Centromeres compete for transmission through meiosis (Malik 2009). Many of these conflicts involve selfish elements that drive (i.e. are transmitted at greater than 50%), particularly via female meiosis (Burt and Trivers 2006). Such selfish genetic elements often impose a cost on fitness, including transposon insertions that render genes nonfunctional, and deleterious alleles that have hitchhiked to high frequency via linkage with a driving centromere. These genomic conflicts also contribute to variation in the number, size and shape of chromosomes in the karyotype (De Villena and Sapienza 2001; Chmátal et al. 2014).

Supernumerary “B” chromosomes, first described a century ago (Wilson 1907), offer a unique system for studying genomic conflicts. B chromosomes are selfish genetic elements that exist alongside the canonical karyotype (or “A” chromosomes) in some individuals of a population. They can vary in number and ploidy and are estimated to occur in at least 15% of eukaryotic species (Burt and Trivers 2006). These selfish chromosomes develop mechanisms to favor their transmission, despite a potential negative impact on organismal fitness (Meiklejohn and Tao 2010).

Sex chromosomes are another focal point for genetic conflicts. Sexually antagonistic selection favors a reduction in recombination on sex chromosomes to increase the association of sexually antagonistic alleles with the sex determining locus (Charlesworth 1991; Charlesworth et al. 2005; Bergero and Charlesworth 2009). Successive events reducing recombination (e.g. inversions) can lead to evolutionary strata with different degrees of differentiation between the sex chromosomes (Bergero and Charlesworth 2009; Zhou et al. 2014). Alternative explanations for the restriction of recombination on sex chromosomes include meiotic drive, heterozygote advantage and genetic drift (Ponnikas et al. 2018).

Sexual conflicts are common and often drive the evolution and turnover of sex chromosomes (Parker 1979; Chapman et al. 2003; Burt and Trivers 2006; Van Doorn and Kirkpatrick 2007; Bachtrog et al. 2014). It has been proposed that female meiotic drive contributes to the evolution of new sex chromosomes via fusions with autosomes, and that karyotype shape affects the types of fusions that occur (Yoshida and Kitano 2012). In fishes and in reptiles, sex chromosome-autosome fusions more often involve Y chromosomes than X, W, or Z chromosomes, which is consistent with them being slightly deleterious and fixed by genetic drift (Pennell et al. 2015; Kirkpatrick 2017).

It has been suggested that some sex chromosomes originated from B chromosomes, or vice versa, based on similarities in their repetitive DNA and transposons, lack of recombination, patterns of heterochromatic gene silencing, and dearth of functional genes (Hackstein et al. 1996; Camacho et al. 2000; Carvalho 2002; Nokkala et al. 2003). However, evidence to support these hypotheses is limited (Charlesworth et al. 2005; Bachtrog 2013; Fraïsse et al. 2017). Definitive tests of these hypotheses will require data from closely related species, as both B chromosomes and young sex chromosomes evolve quite rapidly.

Cichlid fishes have undergone an extraordinary radiation in East Africa, diversifying into more than 1500 species over the last 25 MY (Kocher 2004). Cichlid karyotypes show numerous signs of ongoing genomic conflict. B chromosomes have now been discovered in numerous cichlid species (Poletto et al. 2010; Valente et al. 2014). Many Lake Malawi cichlid species harbor a B chromosome that is present as a single copy and only in females. This B chromosome carries an epistatically dominant female (W) sex-determiner, which likely evolved to promote the transmission of the B chromosome through female meiosis (Clark et al. 2017; Clark and Kocher 2019). In Lake Victoria cichlids, a different B chromosome persists in high frequency (85% of individuals in all species examined). The Lake Victoria B chromosome can be found in one to three copies in males or females, and typically shows no effect on the phenotype (Poletto et al. 2010; Valente et al. 2014). However, in one species of Lake Victoria cichlids, B chromosomes were shown to have an effect on sex determination (Yoshida et al. 2011). Both the Victoria and Malawi B chromosomes have been characterized cytogenetically (Poletto et al. 2010; Fantinatti et al. 2011) and at the sequence level (Clark et al. 2018; Coan and Martins 2018). Similar to other well studied B chromosomes, these cichlid B chromosomes are highly repetitive (Valente et al. 2016). Some autosomally-derived sequence blocks up to ~500kb have become amplified to produce many copies on these B chromosomes. Many of these blocks contain transcribed genes involved in processes related to meiosis and mitosis, suggesting that genes on these B chromosome may be involved in ongoing conflicts to maintain the B chromosomes in the population (Valente et al. 2014; Clark et al. 2018).

African cichlids show an extraordinary diversity of sex chromosomes, and the highest rate of sex chromosome turnover among vertebrates (Gammerdinger and Kocher 2018; Vicoso 2019). Genetic conflicts have contributed to at least some of this diversity. For example, sexual conflict involving color polymorphisms led to the evolution of a novel W sex chromosome (Roberts et al. 2009). Most cichlid sex chromosomes are homomorphic and were therefore identified using sequence markers (Lee et al. 2003; Ezaz et al. 2004; Ser et al. 2010; Lee et al. 2011; Sun et al. 2014). Whole genome sequencing techniques are allowing for the rapid discovery of additional cichlid sex chromosomes (Gammerdinger et al. 2016; Gammerdinger, Conte, Sandkam, Penman, et al. 2018). Several of these novel sex chromosomes involve chromosome fusions (Gammerdinger, Conte, Sandkam, Ziegelbecker, et al. 2018). A chromosome fusion is also associated with an XY sex determination system in the riverine species, *Astatotilapia burtoni* (Roberts et al. 2016). Additionally, polygenic sex determination has been found in *A. burtoni* (Moore and Roberts 2013) and a number of other cichlid lineages (Cnaani et al. 2008; Ser et al. 2010).

The cichlid tribe Oreochromini, comprising ~80-100 species (including the commercially important Nile tilapia, *Oreochromis niloticus*), are unique in having a very large and highly repetitive megachromosome not present in most other cichlids. This megachromosome, referred to here as linkage group 3 (LG3), comprises at least 13.4% of the entire oreochromine genome, and is two to three times larger than any other chromosome in the karyotype (Majumdar and McAndrew 1986; Oliveira and Wright 1998; Ferreira and Martins 2008). LG3 carries a WZ sex-determination locus in several species including the blue tilapia, *O. aureus* (Campos-Ramos et al. 2001; Lee et al. 2004; Conte et al. 2017). The LG3 megachromosome is retained in oreochromines, even when sex determination is controlled by loci on other chromosomes, such as LG1 and LG23 (Eshel et al. 2012; Gammerdinger et al. 2014; Li et al. 2015). A LG3 megachromosome has also been observed in several closely related cichlid lineages including *Pelmatolapia (Tilapia) mariae* (Gammerdinger, Conte, Sandkam, Penman, et al. 2018).

We considered two hypotheses for the origin of the LG3 megachromosome. The first model is that an ancestral autosome acquired a sex determining allele. Following the canonical model of sex chromosome evolution, selection would favor a reduction in recombination (e.g. inversions) that maintain an association between the sex locus and nearby sexually antagonistic variation (Van Doorn and Kirkpatrick 2007). This reduction in recombination would allow an accumulation of deleterious alleles and repetitive elements (Charlesworth et al. 2005; Bachtrog et al. 2014; Vicoso 2019). Under this model, the megachromosome should show conserved synteny with the ancestral autosome, except where gene order has been altered by inversions.

The second model proposed here is that the LG3 megachromosome arose by fusion of an autosome with a highly repetitive B chromosome. Either the autosome or the B chromosome could have been carrying a sex-determination locus prior to fusion. The B chromosome may have carried a sex-determination locus to favor its transmission through meiotic drive (Clark and Kocher 2019). Such a fusion might be favored if it associated sexually antagonistic variation with the sex determiner, or if it contributed to the drive of the B chromosome. Alternatively, the autosome may have carried a sex determination locus and a sexually antagonistic locus on a B chromosome could have favored a fusion of the two chromosomes. In either case, major portions of the megachromosome would show no significant synteny with the ancestral autosome due to the fusion with a B chromosome.

Here we test these hypotheses through a comparative genomic analysis of many cichlid and teleost fish genomes. We present results characterizing the sex megachromosome in the Oreochromini by analyzing synteny, recombination, repeat content, and gene ampliconic arrays. The results shed new light onto the evolutionary origins of this unusual megachromosome and highlight it as a unique sex chromosome for studying genetic conflicts.

## Results

### Cichlid karyotype evolution

The most common teleost karyotype consists of 24 chromosome pairs (2N=48) and is relatively stable within and among lineages (Amores et al. 2014). Previous cytogenetic analyses revealed relative karyotype stability in both Old World cichlids (2N=48) and New World cichlids (2N=44) but identified examples of species-specific fusion and fission events across the family Cichlidae (Poletto et al. 2010). Additional work revealed that African cichlids have experienced two relatively recent chromosome fusions of ancient vertebrate chromosomes (Liu et al. 2013) to create LG7 and LG23 (Conte et al. 2019). In *Astatotilapia burtoni*, a sex chromosome is found on LG5-14, a chromosome fusion that is not found in other cichlid species. The *A. burtoni* genome also experienced another fusion (LG8-24 fused with LG16-21) that has not been associated with sex determination (Ser et al. 2010; Li et al. 2015; Roberts et al. 2016). Figure 1 provides an overview of karyotype evolution in cichlids, as well as the distribution of these fusion events, the LG3 megachromosome and B chromosomes in species from Lake Malawi and Lake Victoria.

**Figure 1.**
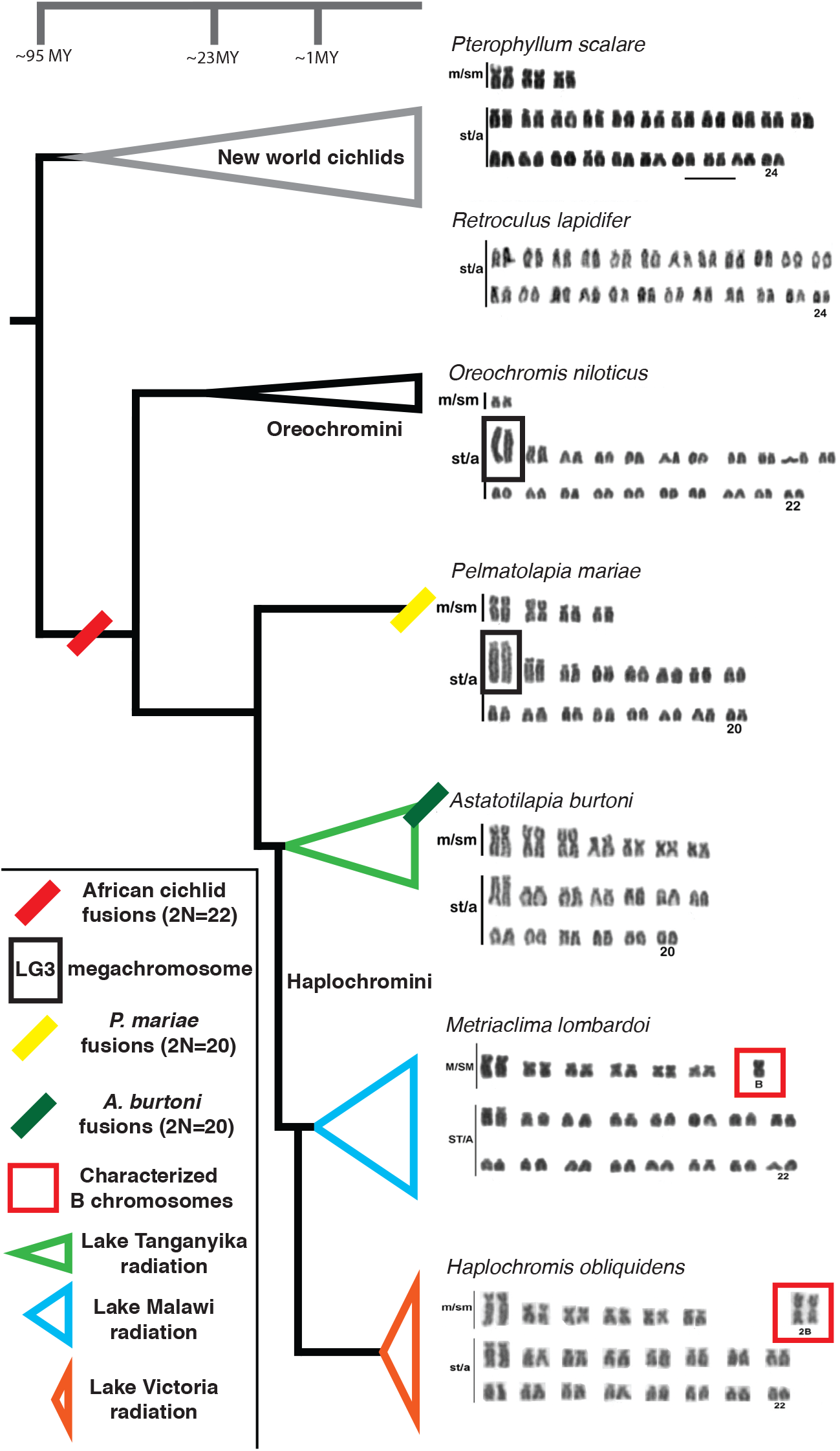
Summary of major karyotype evolution across the phylogeny of cichlids. The Oreochromini and several additional lineages harbor the LG3 megachromosome. The LG3 megachromosome acts as a WZ sex chromosome in at least three different species of *Oreochromis* as well as additional lineages such as *Pelmatolapia mariae*. Karyotypes are adapted with permission (Poletto, Ferreira, Cabral-de-Mello, et al. 2010; Clark et al. 2017).

### WZ sex-determination locus on LG3

Our analysis utilizes two *Oreochromis* genome assemblies – the chromosome-scale assembly of a LG1XX female *O. niloticus* (Conte et al. 2019) and a new chromosome-scale assembly of a LG3ZZ male *O. aureus* (Tao et al. 2020). The LG3 megachromosome was assembled and anchored into 87.6Mbp in the *O. niloticus* assembly. In the *O. aureus* assembly, the size of the LG3 assembly and anchoring is 134.4Mbp. Much of the unanchored sequence in the *O. niloticus* assembly has now been anchored on LG3 in the *O. aureus* assembly (Supplemental File 1).

The *O. niloticus* assembly was previously used to characterize several LG3WZ sex chromosomes. Using the new *O. aureus* assembly as the reference, we re-characterize the sex-determination region on LG3 in *Pelmatolapia mariae* (Gammerdinger, Conte, Sandkam, Penman, et al. 2018) and *O. aureus* (Conte et al. 2017). The *P. mariae* WZ sex-determination locus on LG3 starts at ~25Mbp and extends to 134.4Mbp. The *O. aureus* LG3WZ sex-determination locus starts at ~30Mbp and extends to 134.4Mbp (Supplemental Files 2-3).

### Conservation of synteny

The Japanese medaka (*Oryzias latipes*), provides the most suitable outgroup for studying synteny of LG3 in the Oreochromini. Medaka have a typical teleost karyotype of 24 chromosome pairs and are the most closely related species with high-quality chromosome-scale assemblies (Ichikawa et al. 2017). Due to the fact that the LG3 megachromosome is highly repetitive and contains many gene duplications (Ferreira et al. 2010; Conte et al. 2017), comparison of one-to-one orthologs of five species was necessary to remove alignment artifacts (see Methods). Figure 2 provides a comparison of these five-way one-to-one orthologs of *O. aureus* LG3 to the corresponding medaka chromosome 18. LG3 is divided into three parts (LG3a, LG3a’, and LG3b) based on these patterns of synteny. LG3a consists of the region with conserved synteny comprising the first ~42Mbp of *O. aureus* (99 one-to-one orthologs). LG3a’ consists of the middle ~45Mbp (from ~42Mbp to ~87Mbp) and contains only 12 one-to-one orthologs. LG3b consists of the last 47Mbp of *O. aureus* (from 87Mbp to 134Mbp) and contains zero one-to-one orthologs to medaka. LG3b comprises 35% of the anchored LG3 megachromosome and represents the region potentially originating from a B chromosome fusion. The one-to-one orthologs at the end of medaka chromosome 18 correspond to the final orthologs on LG3a’ in the middle of LG3. The assembly of *O. niloticus* LG3 (87Mbp) shows a similar pattern of synteny to medaka, although the boundary between LG3a’ and LG3b is not as well defined (Supplemental File 4). Several cichlid species outside of the tribe Oreochromini, that do not have the LG3 megachromosome, show conserved synteny to medaka across this entire chromosome (Supplemental Files 5-7). Additionally, it does not appear that LG3a’ and LG3b arose from a different autosome as they do not show detectable synteny with any other chromosomes.

**Figure 2.**
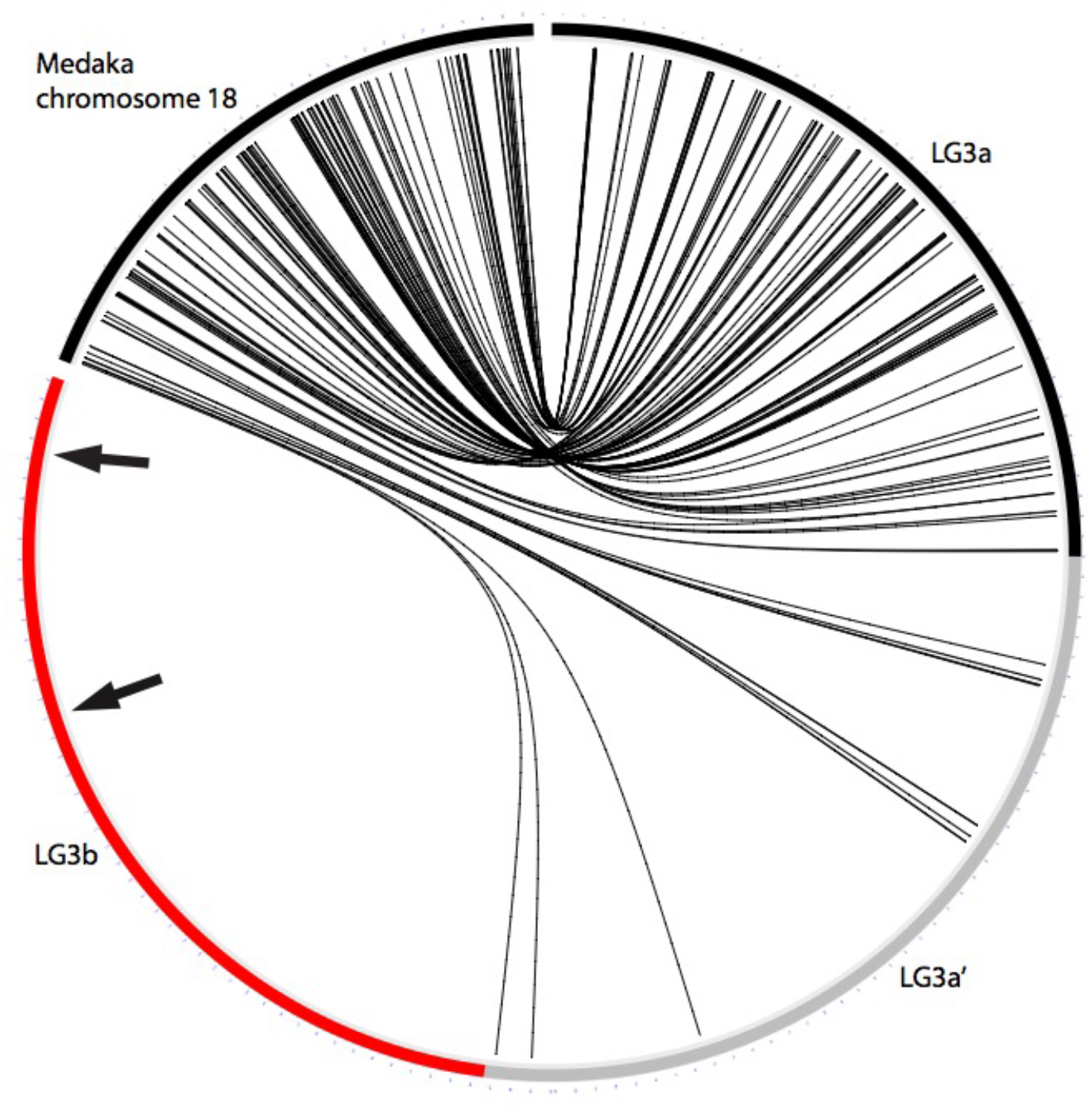
Five-way, one-to-one ortholog alignments of *O. aureus* LG3 to medaka chromosome 18. Interstitial telomeric sequences (ITS) are labeled with black arrows.

Several previous cytogenetic studies have shown that *O. niloticus* LG3 contains two separate interstitial telomere repeat sequences (ITSs) near the center of the chromosome (Chew et al. 2002; Martins et al. 2004). These ITSs may be indicative of chromosome fusion events (Azzalin et al. 2001; Bolzán 2017). Consistent with the cytogenetic studies, the *O. aureus* assembly also contains two telomere repeats arrays (TTAGGG)n that are present on LG3 at 116.9Mbp, 130.6Mbp and at the presumed actual telomeric end at 134Mbp (genome-wide list in Supplemental File 8). The African cichlid specific chromosome fusion events on LG7 and LG23, which occurred before the LG3 megachromosome, have not left traces of ITSs detectable by either cytogenetic studies (Chew et al. 2002; Martins et al. 2004) or the genome assemblies of *O. aureus* and *O. niloticus* (Supplemental File 8).

### Patterns of recombination

The pattern of recombination in *O. niloticus* was previously characterized using a high-density map (Joshi et al. 2018; Conte et al. 2019). LG3a shows the typical sigmoidal pattern of recombination seen on other African cichlid chromosomes, in which recombination rate is low near the telomeres and high in the middle of the chromosome. LG3a’ has lower recombination, and LG3b shows large regions of no recombination (Figure 3). These patterns of recombination also coincide with the patterns of synteny (Figure 2 and Supplemental File 4). LG3a shows a high density of syntenic markers both between *Oreochromis* species, and in comparisons to medaka. LG3a’ shows a lower density of markers and smaller blocks of uninterrupted synteny in both the *O. niloticus* to medaka and *O. aureus* to medaka comparisons. LG3b shows relatively few syntenic markers between oreochromines, and no 1-to-1 orthologs with medaka.

**Figure 3.**
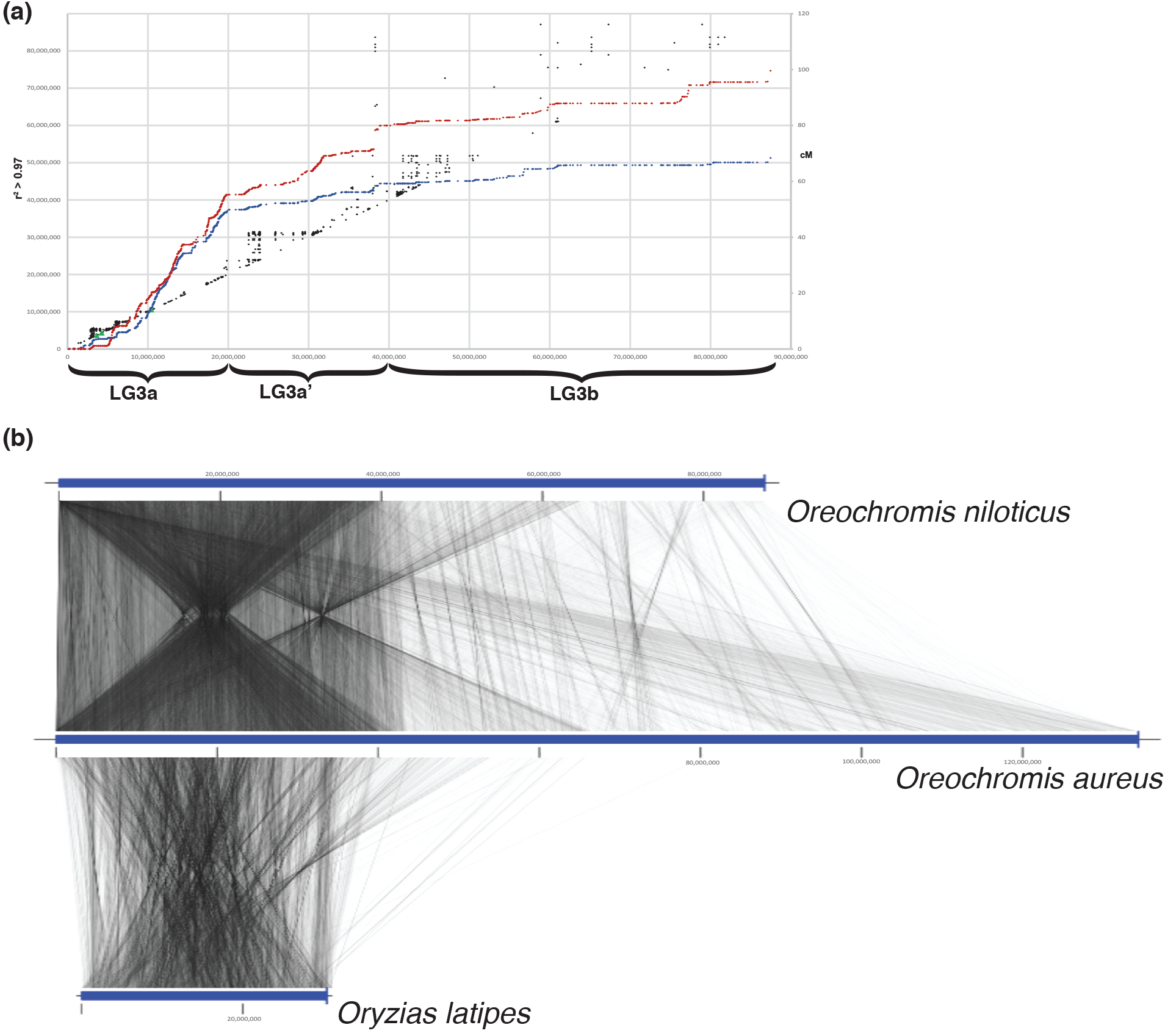
Patterns of recombination in *O. niloticus* correspond to the organization of synteny between *O. niloticus* and *O. aureus* LG3. (a) Recombination of female (red) and male (blue) *O. niloticus* LG3 shown in cM (right) and linkage disequilibrium (r^2^ > 0.97) in black. Adapted with permission from (Conte et al. 2019). (b) Synteny of the 87Mbp assembled and anchored LG3 in *O. niloticus* compared to the 134Mbp assembled and anchored LG3 in *O. aureus* compared to the ancestral chromosome 18 in *O. latipes*.

### Sequence content of the megachromosome

The sequence content of the oreochromine megachromosome is unusual compared to 69 other teleost fish genome assemblies. *O. niloticus* has the highest number of immunoglobulin genes and more than double the number of immunoglobulin transcripts of any other teleost (Supplemental File 9). LG3a’ and LG3b account for 47.4% (100/211) of *O. niloticus* immunoglobulin genes (Supplemental File 10). Subtracting these, *O. niloticus* would have a slightly above average count (111 versus the teleost average of 101). The Oreochromini also have the largest amount of total sequence of any teleost annotated as endogenous retrovirus (ERVs), of which LG3a’ and LG3b account for 13.8% (1.06Mbp of the total 7.67Mbp genome-wide). However, the Oreochromini do not have the highest number of ERV insertion events. This either suggests a fragmented and incomplete representation of these elements in teleost assemblies constructed from short-read sequence data (Conte and Kocher 2015) and/or that oreochromine ERVs are more recent and intact, resulting in fewer annotated ERV copies than more highly decayed ERVs in other species. The Oreochromini also have the highest number of annotated long non-coding RNAs (lncRNAs) among teleosts. LG3a’ and LG3b account for 13.1% of these lncRNAs. Additionally, LG3b has a high density of zinc-finger proteins relative to the rest of the genome, although the overall number of these zinc-finger proteins is similar to that in other teleosts. Finally, a gene ontology (GO) enrichment analysis of LG3b identified several significantly enriched terms, all related to immune regulation and immune response (Supplemental Files 11-12).

The megachromosome contains several large, highly repetitive, ampliconic gene arrays which are commonly found on both sex chromosomes and B chromosomes of other species (Bellott et al. 2010). The extent of these ampliconic arrays can be seen on a chromosome scale by examining sequence similarity across LG3 (Supplemental File 13). These ampliconic gene expansions are found most frequently in the non-recombining regions of LG3b. However, some of these genes have also expanded throughout LG3 and are also seen in lower copy numbers in the freely recombining region on LG3a and lower recombining region of LG3a’. A table of genes that have undergone expansion on LG3 is provided in Supplemental File 14.

### Patterns of transposable elements on the megachromosome

The LG3 megachromosome is distinguished by having the highest density of repetitive elements across the genome (Ferreira et al. 2010; Conte et al. 2019), which may be a signature of a fusion with a B chromosome. B chromosomes in cichlids have been characterized as having a much higher content of specific TE families relative to the A genome (Coan and Martins 2018). One explanation for this might be that B chromosomes can act as a “safe-haven” for particular TEs (McAllister and Werren 1997; Camacho et al. 2000; Werren 2011). Therefore, B chromosomes may be more likely to contain TE families diverged from copies on the A. In the most extreme case, one might also expect selfish B chromosomes to contain private TE families not found in the A chromosomes. *O. aureus* LG3 contains three different unknown TE families that were not found on any other chromosome and which are present in at least 100 copies (see Methods), defined here as “completely private TE families”. Additionally, *O. aureus* LG3 contains six additional TE families that were present in at least 100 copies and were almost exclusively found on LG3 only (>98% of copies), defined here as “predominately private TE families” (Supplemental File 15). One of these families was annotated as a DNA/Dada element while the remainder were unknown elements. These private TE families on LG3 were mostly found on LG3a’ and LG3b, while very few copies of these TE families were found on LG3a. The rest of the *O. aureus* genome contains only two chromosomes (LG4 and LG13) with completely private TE families (one each) and no other chromosomes containing a predominately private TE family. The private TE results are similar for *O. niloticus* LG3 compared to the rest of the genome (Supplemental File 15).

The age of these private TEs is an important factor to consider as well. For example, if the private TEs were all very recent in age, then perhaps they arrived well after the potential B chromosome fusion event. On the other hand, if the private TEs were older in age, then this may be evidence that they evolved on the original B chromosome prior to a potential fusion. The *O. aureus* repeat landscape (Supplemental File 16) is similar to the *O. niloticus* repeat landscape (Conte et al. 2017; Conte et al. 2019). The completely private TE copies share a similar shape to the whole genome with copies of all ages as is the case for the predominately private TE copies (Supplemental File 16).

## Discussion

Several previous studies have noted similarities between B chromosomes and sex chromosomes and suggested that they may have shared origins (Camacho et al. 2000; Carvalho 2002). Due to the fact that African cichlids have several well-characterized B chromosomes and a high rate of sex chromosome turnover, we sought to test two separate hypotheses concerning the origin of a prominent sex megachromosome in the Oreochromini. The first hypothesis is that an ancestral autosome gained a new sex determining allele, upon which sexually antagonistic selection favored a reduction in recombination via a series of inversions. These reductions in recombination then allowed for the accumulation of many repetitive sequences, eventually resulting in the present megachromosome. The second hypothesis is that an ancestral autosome fused with a highly repetitive B chromosome, where either the B chromosome harbored a dominant sex-determination locus or where the ancestral autosome carrying a sex-determination locus fused with a B chromosome to resolve a conflict. In either case, under this model much of the repetitive nature of the LG3 megachromosome is derived from the initial B chromosome that was later incorporated into the A genome via the fusion. This may have coincided with the spread of heterochromatin to silence transposable elements on the B chromosome and a reduction of recombination outward from the fusion.

The LG3 megachromosome functions as a WZ sex chromosome in several known tilapia species. We have characterized the sex interval as ~25Mbp-134.4Mbp in *T. mariae* and ~30Mbp-134.4Mbp in *O. aureus* using the new *O. aureus* ZZ reference. If the LG3 megachromosome arose as a conventional sex chromosome, we would expect it to contain orthologous sequences across the entire length of the chromosome. We would also expect fewer orthologous sequences in the region of reduced recombination that has accumulated repetitive sequence. Alternatively, if the megachromosome arose via fusion with a B chromosome, we would expect there to be a large region with no orthologous sequences compared to the ancestral autosome. Therefore, the primary test of these hypotheses was to examine the syntenic relationships of the LG3 megachromosome with an example of the ancestral autosome. The medaka genome assembly was chosen for this synteny comparison since it shares the 24 chromosomes common to a majority of teleosts (Guyomard et al. 2012; Amores et al. 2014) and is representative of an ancestral karyotype that has not undergone many rearrangements (Kasahara et al. 2007). Additionally, several recent medaka genome assemblies (Ichikawa et al. 2017) are also very accurate and complete, which benefited our analysis. Figure 2 shows the alignment of the one-to-one orthologs between *O. aureus* LG3 and the corresponding medaka chromosome 18. The majority of one-to-one orthologs are found in LG3a, and a few are present in LG3a’. LG3b contains zero orthologs across at least 47Mbp. The 47Mbp comprising LG3b alone is larger than all but one other cichlid chromosome (the African cichlid-specific fusion of LG7 is the only larger chromosome at 66Mbp). This finding is consistent with the B chromosome fusion hypothesis. Considering the synteny results in the context of the first hypothesis, it is difficult to imagine a series of inversions and/or rearrangements that could have resulted in such a large portion of the chromosome that does not contain any ancestral orthologs.

The second test of the hypotheses was to examine a notable characteristic of fused chromosomes: the presence of interstitial telomere repeat sequences (ITSs) (Bolzán 2017). ITSs have been shown to be markers of ancient chromosome fusion events in several well-studied vertebrate genomes (Ijdo et al. 1991; Azzalin et al. 2001; Tsipouri et al. 2008). ITSs were found at two places on LG3b in *O. aureus* (116.9Mbp and 130.6Mbp) and are not found on any other chromosomes. Previous cytogenetic studies in *O. niloticus* discovered two ITSs in roughly the middle of the long arm of LG3 (Chew et al. 2002; Harvey et al. 2003; Martins et al. 2004). So, the placement of the *O. aureus* ITSs is not completely consistent with the *O. niloticus* cytogenetic studies. This discrepancy may be explained either by differences in the structure of LG3 between these two Oreochromini species and/or limitations in the accuracy of the assemblies. The fact that two distinct ITSs are found on LG3b raises the possibility of multiple fusion events, inversion event(s) on LG3, and/or a more complicated history than either of our two hypotheses account for. There are also several important caveats that suggest the presence of these ITS sequences on LG3 may not be due to a chromosome fusion event. It is possible that ITSs were carried by TE families specific to subtelomeric regions or could have been inserted due to telomerase-mediated repair of double-stranded breaks (Bolzán 2017). Interestingly, it has been shown that a large portion of the LG3 megachromosome does not pair during the early pachytene stage of meiosis (Foresti and Oliveira 1993; Ocalewicz et al. 2009). ITSs can play a variety of roles in the organization and stability of chromosomes (Aksenova and Mirkin 2019), and the ITSs on LG3 may be involved in meiotic instability. It is possible that the absence of pairing of the megachromosome during meiosis may be a relic of its origin from a fused B chromosome and the ITSs may be part of the reason for the pairing abnormality. Nonetheless, the presence of these ITSs on LG3b is more consistent with the B chromosome fusion hypothesis.

If the megachromosome had originally evolved as a B chromosome for some time before a potential fusion and incorporation into the A genome, one might expect it to contain copies of unique TE families that are not present in the A genome. Indeed, these private TE families are more common on LG3 relative the other chromosomes (Supplemental File 15). These private TE families are mostly located on LG3a’ and LG3b. However, this is also where recombination is lower and so the efficacy of purging deleterious TE indels is also lower. The wide range in the age of the private TEs suggest that perhaps they evolved on a former B chromosome and not more recently (Supplemental File 16). Although these private TE families are much more common on the LG3 megachromosome, the fact that two private TE families also occur on other chromosomes may mean that private TE families can arise in different ways and may not be diagnostic of a B chromosome. This piece of evidence does not strongly support either hypothesis.

The remainder of the results gathered describing the LG3 megachromosome including recombination, gene content, and the ampliconic regions are not able to distinguish between the two hypotheses regarding the origin of the megachromosome since each feature shows similar patterns in both B chromosomes and sex chromosomes (Camacho et al. 2000). The synteny, ITSs, and private TE results that characterize the megachromosome favor rejection of the canonical sex chromosome hypothesis to various degrees, but do not provide overwhelming evidence to reject this hypothesis at this point. Table 1 provides a summary of the results and how well they support each hypothesis or both.

**Table 1.**
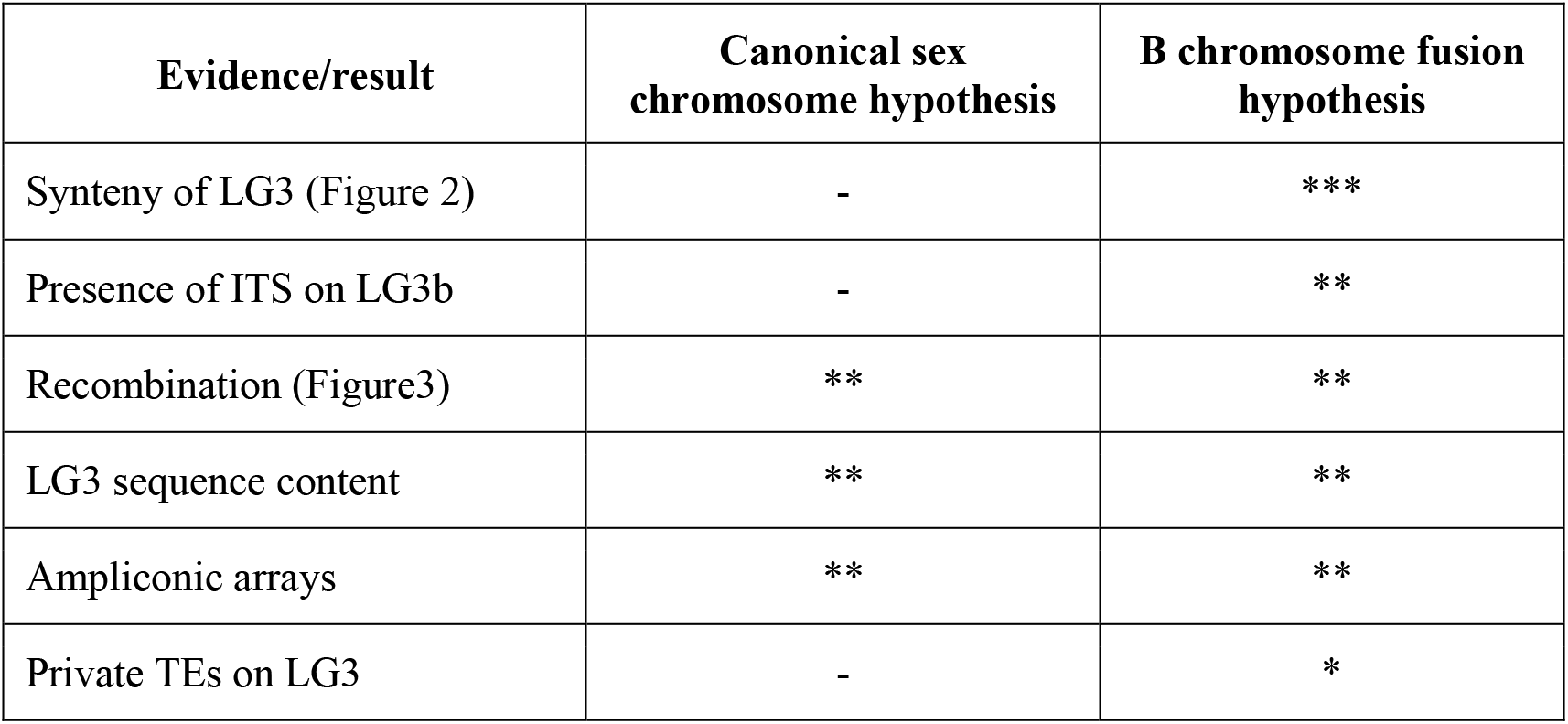
Summary table of whether each result supports one of two hypotheses regarding the origin of the LG3 megachromosome or if the result supports both hypotheses. *** indicates strong support, ** indicates support, but with caveats, * indicates some support, - indicates no support.

While the oreochromine sex megachromosome provides a very intriguing new case to test the B chromosome fusion sex chromosome hypothesis, data from more cichlid species with and without B chromosomes are needed. Fusions of B chromosomes with A chromosomes are not unprecedented. The fusion of a B chromosome with an autosome has been shown in the fungus *Nectria haematococca*, where the B chromosome provides antibiotic resistance (Miao et al. 1991). Another B chromosome fusion has been shown in the grasshopper *Eyprepocnemis plorans*, where B chromosomes interact with nucleolar organization regions of A chromosomes that result in polymorphic fusion events (Teruel et al. 2009). B chromosomes that physically interact with A chromosomes may be more predisposed to fusions. Yet another fusion has been reported in laboratory stock of the medfly *Ceratitis capitata*, where small B chromosomes fused with the X chromosome, creating polymorphism in X chromosome size (Basso and Lifschitz 1995). A situation similar to this polymorphism, where B chromosomes are fused in some individuals of a population but not others, may provide the most suitable situation for studying the questions of B chromosome and sex chromosome origin. It remains to be seen if such a situation still exists in any Oreochromini species or perhaps in other cichlid groups.

The LG3 megachromosome acts as a WZ sex chromosome in some species (e.g. *O. aureus* and *P. mariae)*, but not in others (e.g. *O. niloticus)*. There is not yet any evidence of heteromorphism between the W and Z chromosomes in any of these species, which is not unusual for cichlid sex chromosomes. These species may have recently fixed either the W or Z chromosome, perhaps as the result of the frequent turnover of sex chromosome systems in this lineage. It will be easier to reconstruct the evolutionary history of these sex chromosome turnovers once the sex-determining gene(s) on LG3 are identified. It may appear that *O. niloticus* and *O. aureus* LG3 differ dramatically given the size difference (Figure 3). However, it is important to point out that the *O. niloticus* assembly contains more unanchored sequences than the new *O. aureus* assembly. Many of these unanchored sequences in *O. niloticus* assembly contain the LG3W sex-determination interval (Conte et al. 2017).

It should also be noted that megachromosomes do not appear to have evolved in other cichlid lineages. Most of the sex chromosomes identified in East African cichlids have evolved quite recently and show modest levels of differentiation. The extreme differentiation of LG3 in oreochromines compared to other cichlids suggests that this megachromosome is much older than other cichlid sex chromosomes. There is no evidence to suggest that these more recent sex chromosomes are on a trajectory to become megachromosomes.

Comparisons across many teleost genomes indicate that the LG3 megachromosome has a distinct repetitive sequence content (Supplemental Files 9 and 10). What process(es) may have caused this chromosome to acquire so many repeated sequences? Molecular evolutionary arms races are known to play a role in genome size evolution (Ågren and Wright 2015; Kapusta et al. 2017; Cosby et al. 2019). An arms race in which zinc-finger proteins evolve to bind to and suppress transcription of endogenous retroviruses (ERVs) has been well documented in mammalian genomes (Bruno et al. 2019; Wolf et al. 2020). Oreochromines have an average amount of zinc-finger proteins compared to other teleosts, but LG3 and particularly LG3b contains a large fraction of the zinc-finger proteins in their genome (Supplemental File 10). So, it is possible that the increased number of zinc-finger proteins on LG3 are involved in silencing ERVs and perhaps other transposable elements. While there is ongoing conflict between host and ERVs, studies have shown that ERVs can also be co-opted and contribute to host immunity and antiviral defense (Lynch 2016; Chuong et al. 2017; Frank and Feschotte 2017). It is possible that a by-product of this arms race is a benefit to the host immune system. This could explain why this megachromosome is still present in species where it is not the sex-determining chromosome. lncRNAs also play a role in the immune response to viral infection (Satpathy and Chang 2015; Ouyang et al. 2016) and are very abundant on LG3b and compared to other teleost genomes. These lncRNAs may also be intertwined in a complicated arms race on this megachromosome.

High-quality genome assemblies of sex chromosomes are becoming more available (Liu et al. 2019; Peichel et al. 2019). An assembly of the neo-Y chromosome of *Drosophila miranda* showed that the neo-Y chromosome has expanded and many of the genes have been amplified to high copy number (Bachtrog et al. 2019). Many of the genes on this neo-Y have functions related to chromosome organization such as chromosome organization/segregation, mitotic cell cycle, meiosis, spindle assembly, and kinetochore assembly. This gene content is very similar to the gene content of the two well described African cichlid B chromosomes in Lake Victoria (Valente et al. 2014) and Lake Malawi (Clark et al. 2018). The *D. miranda* neo-Y study did not examine the potential involvement of a B chromosome, but the similarities are striking. Additional high-quality sex chromosomes will continue to become more available and it will be interesting to see what insights they will provide for investigating the similarities and origins of B chromosomes and sex chromosomes.

## Conclusion

This study presents a new case study to address questions about the possible origin of sex chromosomes from B chromosomes. We speculate that the sex megachromosome in oreochromine cichlids arose via the fusion of an autosome with a B chromosome to resolve a conflict of a W sex determination locus. Our work documents the structure of an extreme sex chromosome in African cichlids and provides a benchmark against which the characterization of sex chromosomes in this group can be compared. More generally, it provides a new system for studying the evolutionary arms races that shape chromosome architecture.

## Materials and Methods

### WZ sex-determination locus on LG3

Whole genome pooled sequencing of *T. mariae* males (SRR6660983/SRR6660984) and females (SRR6660979/ SRR6660980) and *O. aureus* males (SRR5121056) and females (SRR5121055) were aligned to the *O. aureus* ZZ reference assembly using BWA mem version 0.7.12-r1044 (Li 2013). Picard version 2.1.0 was used to sort the SAM files, mark duplicates, and index the BAM files. Methods to calculate F_ST_, and determine WZ sex-patterned SNPs, were previously described (Gammerdinger, Conte, Sandkam, Penman, et al. 2018) and used here.

### Synteny analysis

Five-species one-to-one orthologs were computed using OrthoFinder version 2.3.3 with the ‘-I 5 - S diamond’ options enabled. NCBI RefSeq protein annotations from five fish species were used as input for this analysis. They include the *O. niloticus* (GCF_001858045.2), *Archocentrus centrarchus* (GCF_007364275.1), *Astatotilapia calliptera* (GCF_900246225.1), *Metriaclima zebra* (GCF_000238955.4), and the outgroup Japanese medaka *Oryzias latipes* (GCF_002234675.1). OmicCircos (Hu et al. 2014) version 1.16.0 used with R version 3.4.1 was used to generate the plots of these ortholog synteny comparisons.

### Whole chromosome synteny analysis with nucmer and genoPlotR

MUMmer version 4.0.0.beta2 (Kurtz et al. 2004) was used for whole genome synteny analysis. First, the ‘nucmer’ program was used to generate all-by-all comparisons of nucleotide sequences. The ‘delta-filter’ program was used to filter these alignments with the following options: *-1 -l 50 -i 50 -u 50*. Finally, the ‘show-coords’ program was used to convert the ‘delta-filter’ output into a tab-delimited file with the following options: *-I 50 -L 50 -l -T -H*. The alignments were visualized (Figure 3) using the R package genoPlotR version 0.8.9 (Guy et al. 2010).

### Analysis of *Oreochromis* LG3b content and comparison across teleost genomes

We downloaded 69 publicly available teleost genomes (listed in Supplemental File 9) that have RefSeq annotation available from the NCBI FTP server (Anon.). Immunoglobulin genes and lncRNAs were extracted from the RefSeq annotation GFF file which was also downloaded from the FTP server. These correspond to annotations that were current as of RefSeq release 94. RepeatModeler (Smit, AFA, Hubley 2010) (*version open-1.0.8*) was used to identify and classify repeats for each of the 69 teleost genomes, separately. These *de novo* repeats specific to each teleost genome were combined with the RepBase-derived RepeatMasker libraries (Bao et al. 2015), separately. RepeatMasker (Smit, AFA, Hubley, R & Green 2010) (*version open-4.0.5*) was run then on each assembly using NCBI BLAST+ (*version 2.3.0+*) as the engine (‘*-e ncbi*’) and specifying the combined repeat library (‘*-lib*’). The more sensitive slow search mode (‘*-s*’) was used.

### Gene Ontology enrichment analysis

There is currently no Gene Ontology annotation of the most recent *O. niloticus* genome assembly (O_niloticus_UMDNMBU/ GCA_001858045.3). Therefore, BLASTX (version 2.2.28+) was used to align *O. niloticus* NCBI RefSeq transcripts (O’Leary et al. 2016) against Swiss-Prot (Bateman 2019) ‘release-2019_01’ which had been formatted using the Trinotate (version 3.1.0) (Bryant et al. 2017) ‘Build_Trinotate_Boilerplate_SQLite_db.pl Trinotate’ command. Those transcripts on LG3b which had a significant BLASTX hit were compiled into a list of 614 gene symbols. This list was uploaded as an ID List for the PANTHER Gene List Analysis (Thomas et al. 2003) using *Homo sapiens* as an organism and performing a statistical overrepresentation test with the GO-Slim Biological Process annotation set. A Fisher’s exact test was performed, and the false discovery rate (FDR) was calculated. Only significant results with a FDR of P < 0.05 were kept.

### Dotplot of *O. aureus* LG03

Gepard version 1.4 (Krumsiek et al. 2007) was used to align *O. aureus* LG03 to itself. A word length of 100bp and a window size of 0bp were used.

### Telomere repeat analysis

Telomere repeat sequences were detected via RepeatMasker output which was run on the *O. aureus* assembly in the same way as described above. They telomere repeat annotations were filtered based on repeat length. For *O. aureus*, this repeat length was set at 100 or greater.

## Acknowledgments and funding information

We acknowledge all members of the UMD cichlid labs for their countless discussions on this project. We acknowledge the University of Maryland supercomputing resources (www.it.umd.edu/hpcc/) made available in conducting the research reported in this paper. This study was supported by funding from the US National Science Foundation (DEB-1830753 to TDK).

