## Supplemental Information for "Evolution of a sex megachromosome"

Supplemental File 1 – Comparison anchored chromosomes of the *O. niloticus* genome assembly (GCF_001858045.2) and the *O. aureus* genome assembly.

Supplemental File 2 – Male versus female $F_{ST}$ for both *T. mariae* and *O. aureus* using the *O. aureus* ZZ genome as a reference.

Supplemental File 3 – Allele frequency of W-patterned SNPs on LG3 for *T. mariae* and *O. aureus* using the *O. aureus* ZZ genome as a reference.

Supplemental File 4 – Comparison of five-species, one-to-one orthologs of *O. niloticus* LG03 (NC_031967.2) and *O. latipes* chromosome 18 (NC_019876.2).

Supplemental File 5 – Comparison of five-species, one-to-one orthologs of *O. niloticus* LG03 (NC_031967.2) and *M. zebra* lg3 (NC_036782.1).

Supplemental File 6 – Comparison of five-species, one-to-one orthologs of *A. calliptera* chromosome 3 (NC_039304.1) and *O. latipes* chromosome 18 (NC_019876.2).

Supplemental File 7 – Comparison of five-species, one-to-one orthologs of *A. centrarchus* chromosome 18 (NC_044363.1)/ chromosome 18 unlocalized scaffold 23 ctg 1 (NW_022060145.1) and *O. latipes* chromosome 18 (NC_019876.2).

Supplemental File 8 – Location of telomere repeat sequence (TTAGGG)n in the *O. aureus* assembly.

Supplemental File 9 – Comparison of 69 teleost genomes available in NCBI RefSeq. Comparison includes ERVs, lncRNAs, and immunoglobulin genes. The maximum value for each column is highlighted in yellow.

Supplemental File 10 – Comparison across chromosomes of ERVs, LINE/L2 elements, LTR/Gypsy elements, LINE/Rex-Babar elements, lncRNAs, immunoglobulin genes/transcripts, zinc-finger proteins, and genes for both the *O. niloticus* and *O. aureus* assemblies.

Supplemental File 11 – Gene enrichment overrepresentation pie chart results.

Supplemental File 12 – Overrepresentation test of the genes on LG3b and the seven significant terms.

Supplemental File 13 – Dotplot of *O. aureus* LG03 compared to itself to highlight highly repetitive regions across the chromosome.

Supplemental File 14 – The copy number of each of the ampliconic genes on *O. niloticus* LG03.

Supplemental File 15 – Proportion and number of private TE families for each chromosome in the *O. aureus* and *O. niloticus* assemblies.

Supplemental File 16 – a) Repeat landscape of the entire *O. aureus* genome assembly, b) repeat landscape of the completely private TEs on LG3, c) repeat landscape of the predominately private TEs on LG3, and d) repeat landscape of both the completely and predominately private TEs on LG3, combined.
