## Supplementary figures and images for "Evolution of a sex megachromosome"

### Supplemental File 2

*T. mariae* males vs females $F_{ST}$

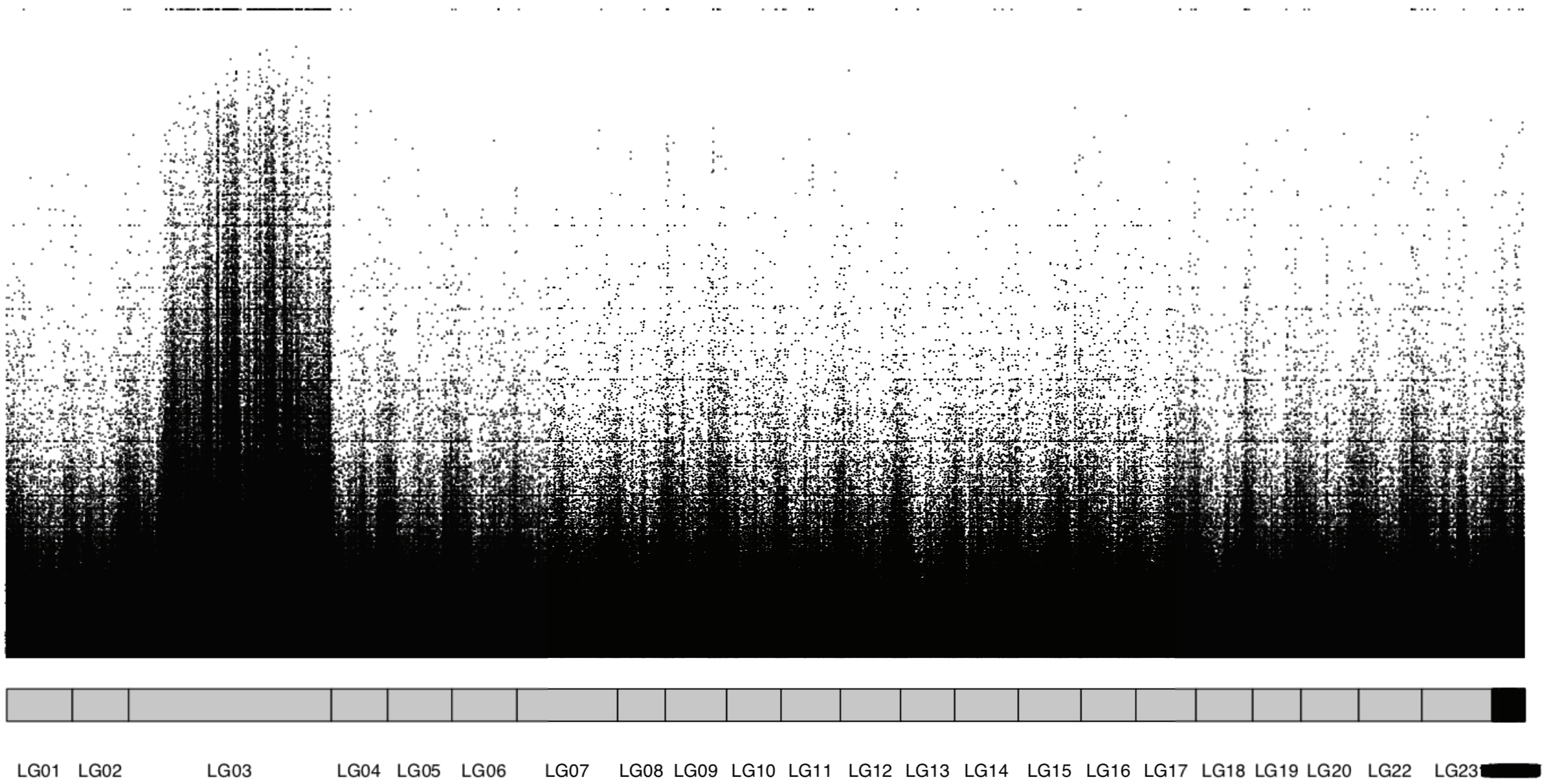


*O. aureus* males vs females $F_{ST}$

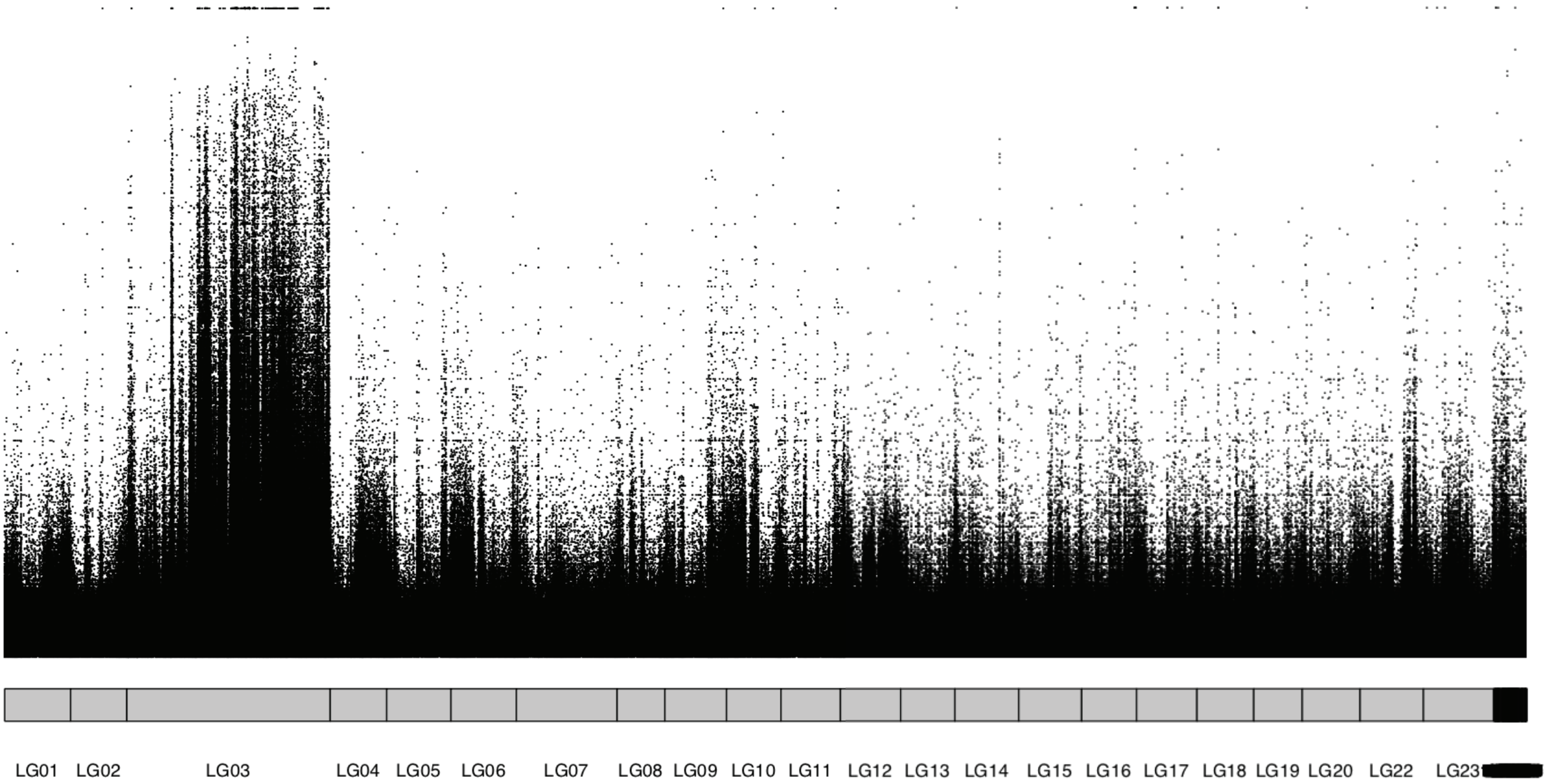

### Supplemental File 3

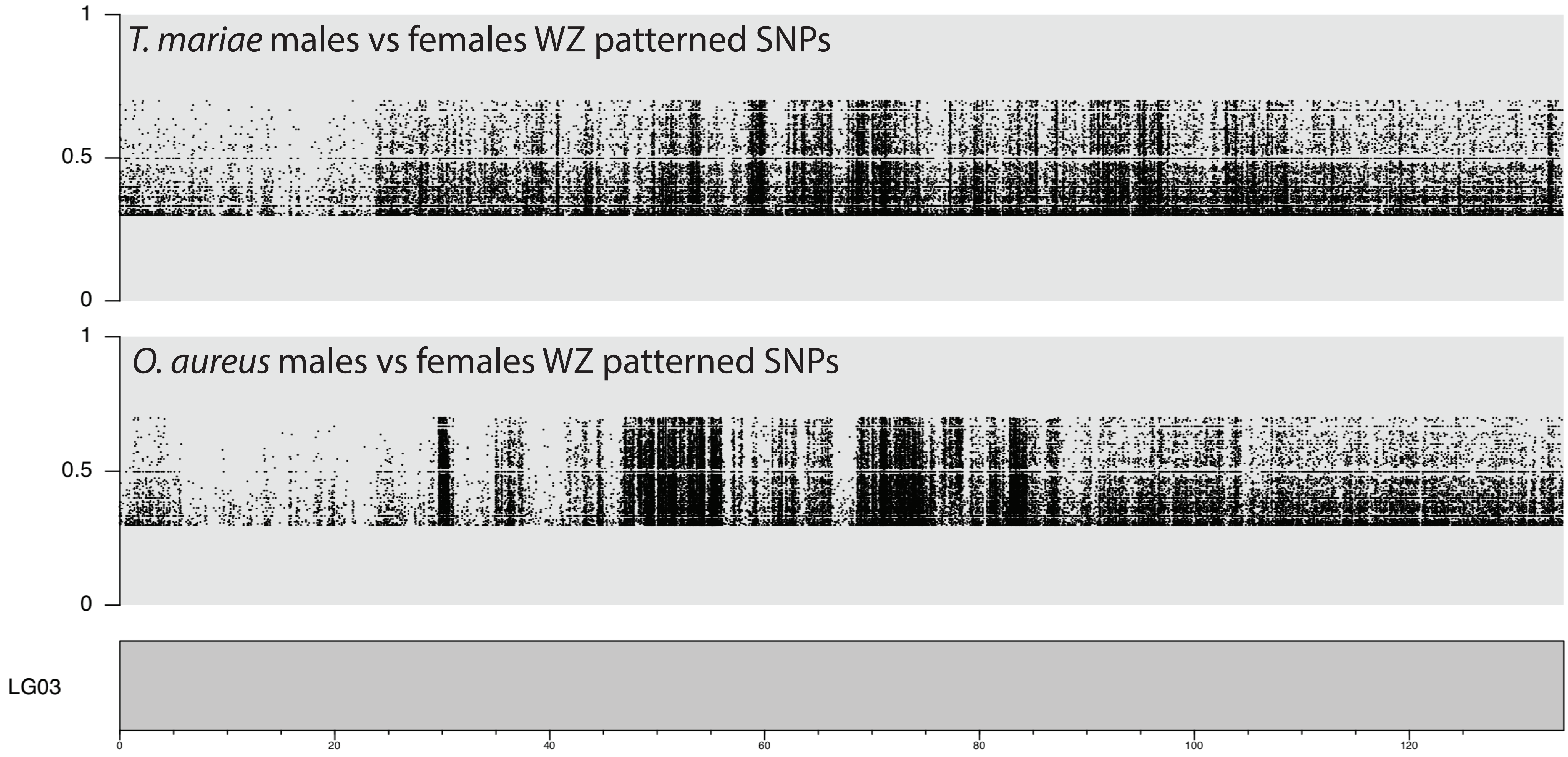

### Supplemental File 4

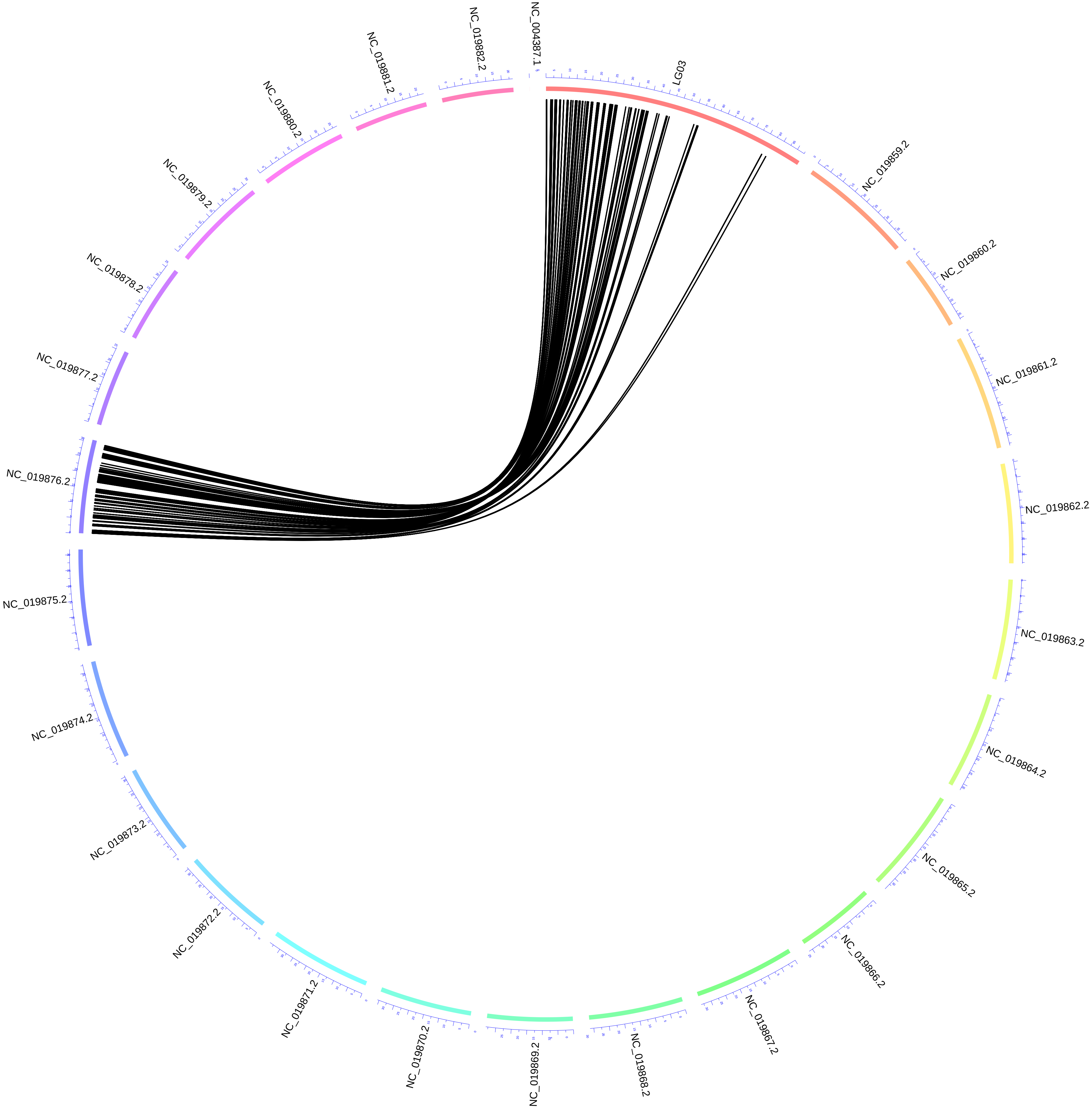

### Supplemental File 5

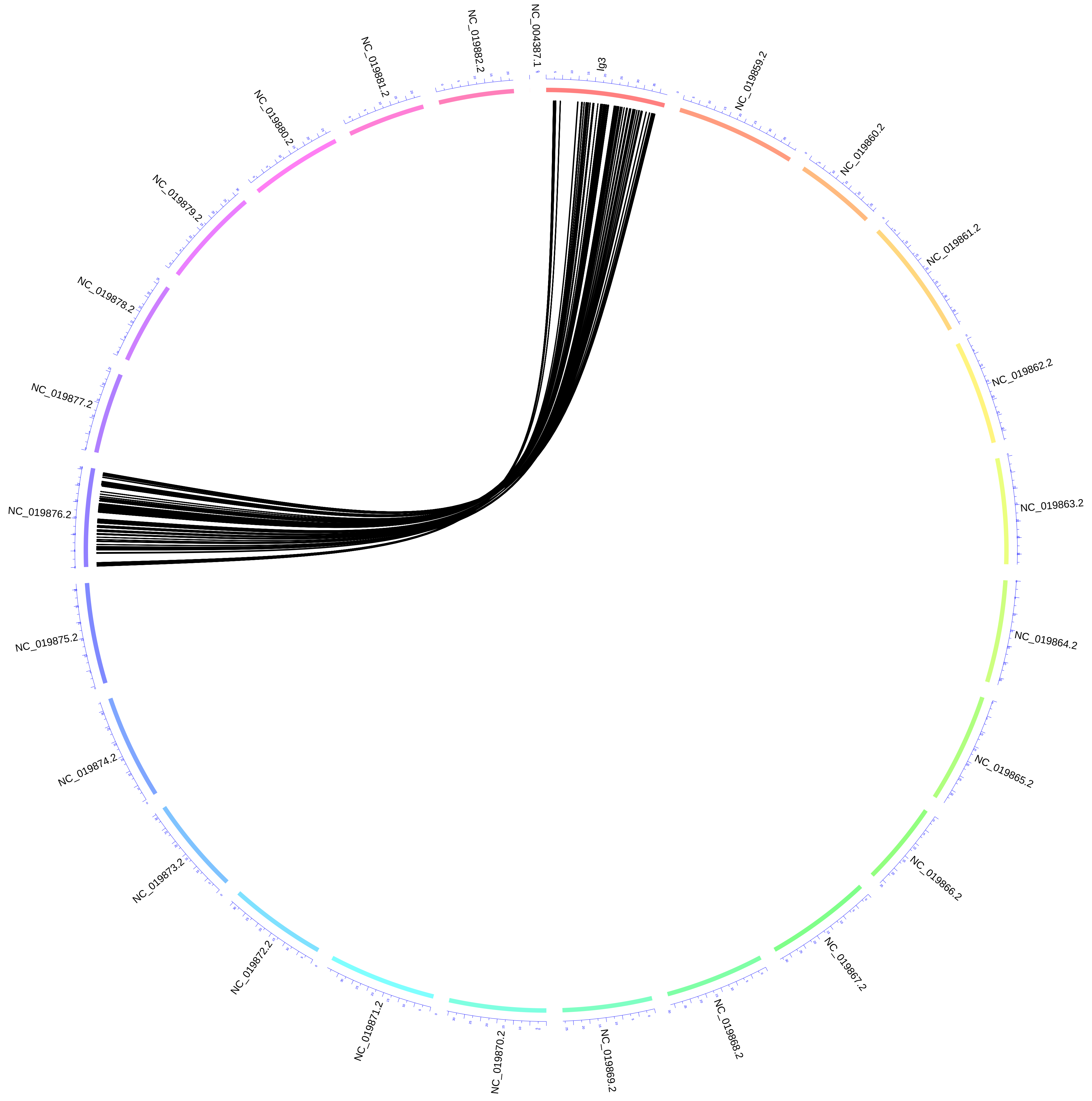

### Supplemental File 6

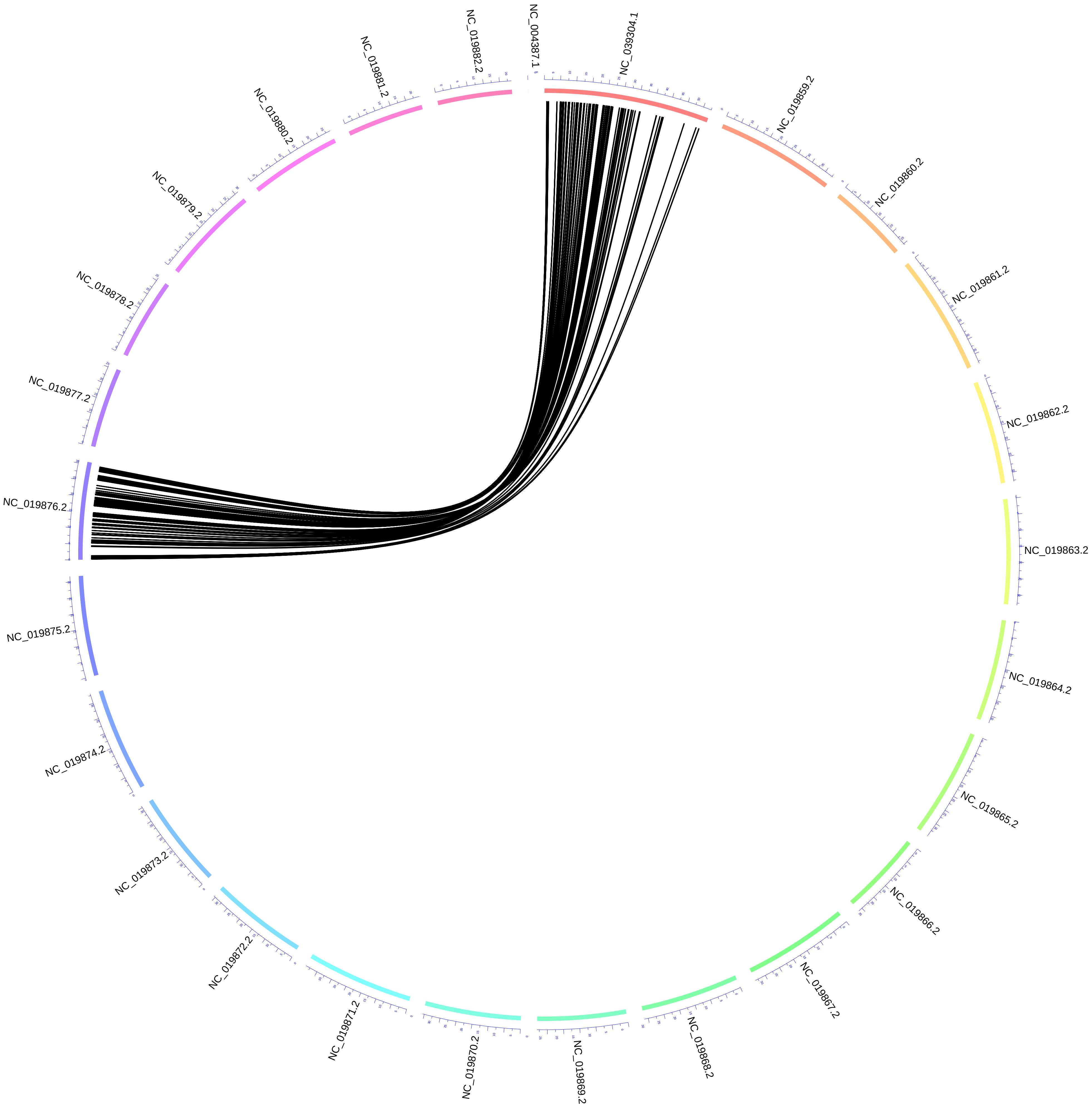

### Supplemental File 7

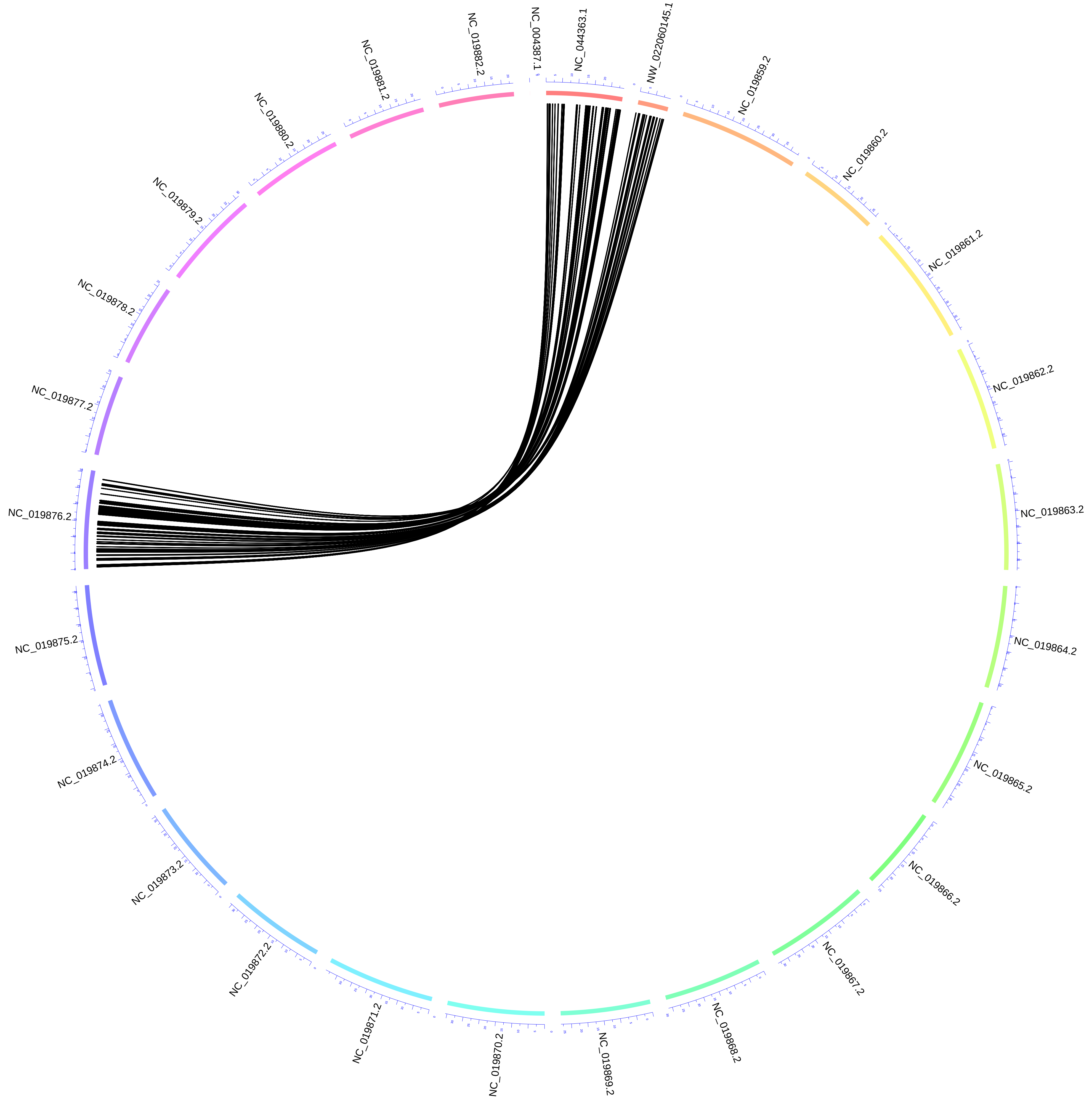

### Supplemental File 11

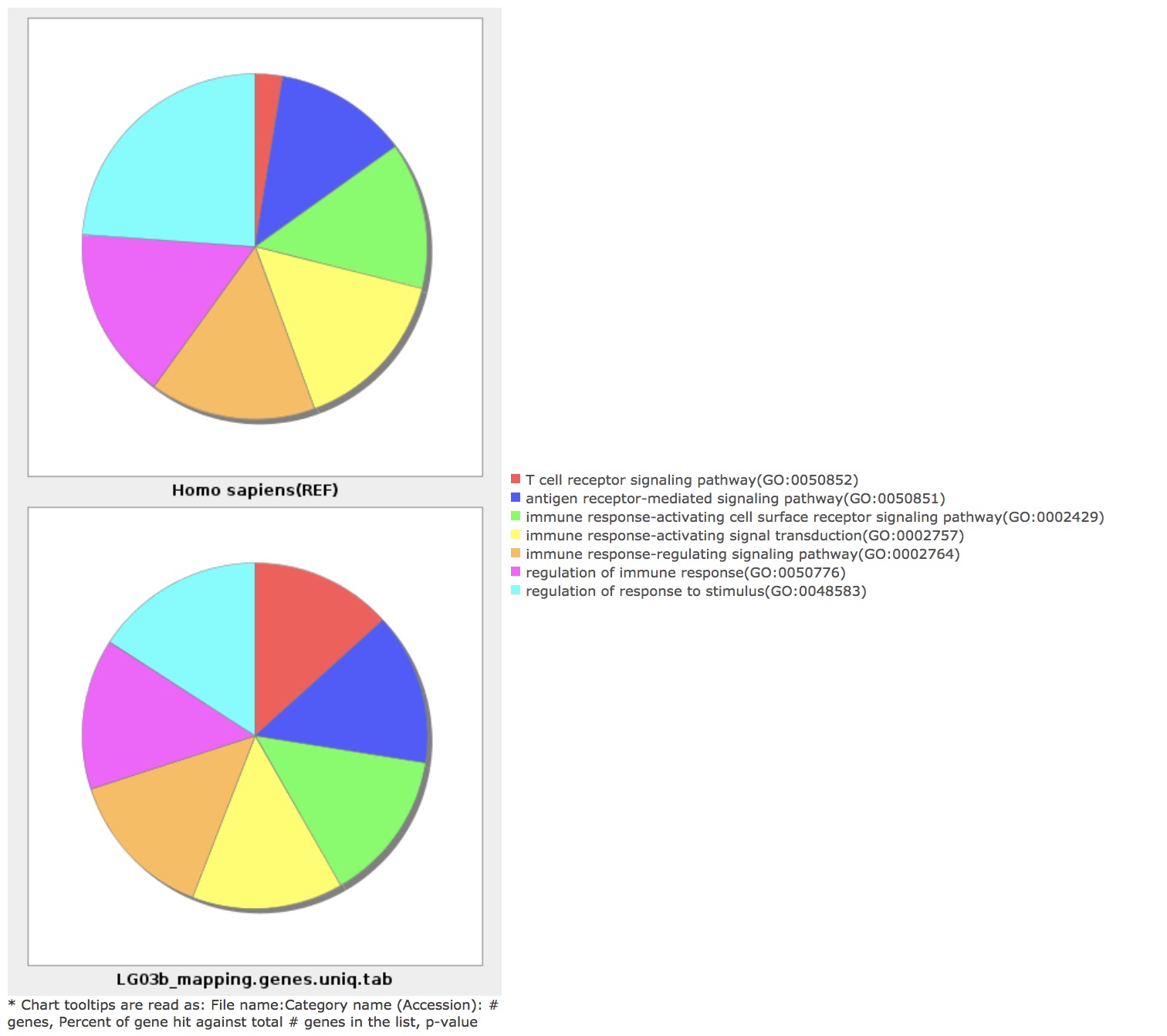

### Supplemental File 13

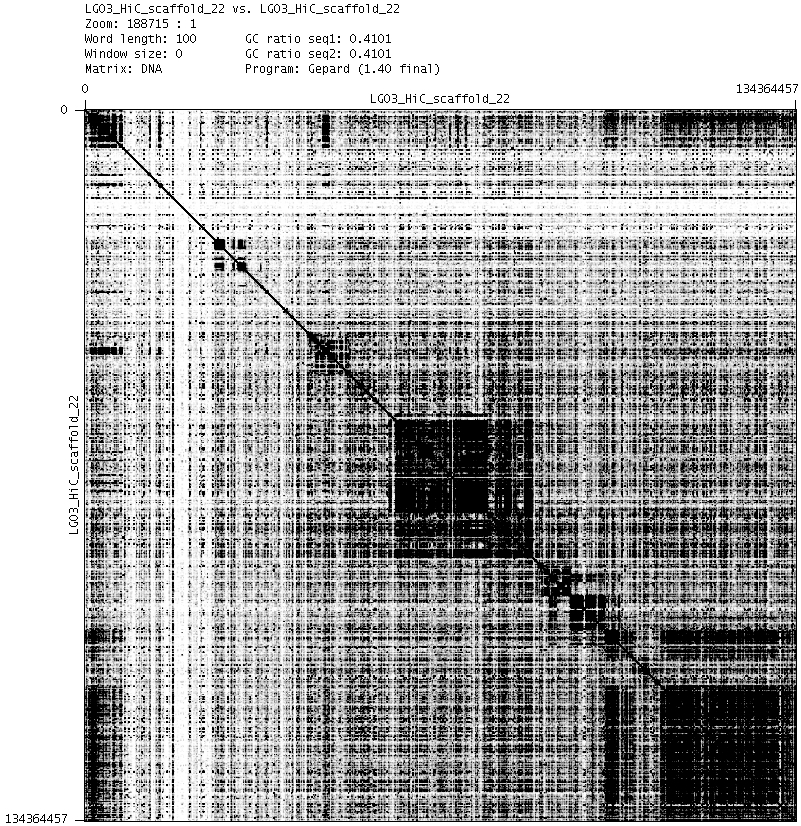

### Supplemental File 16

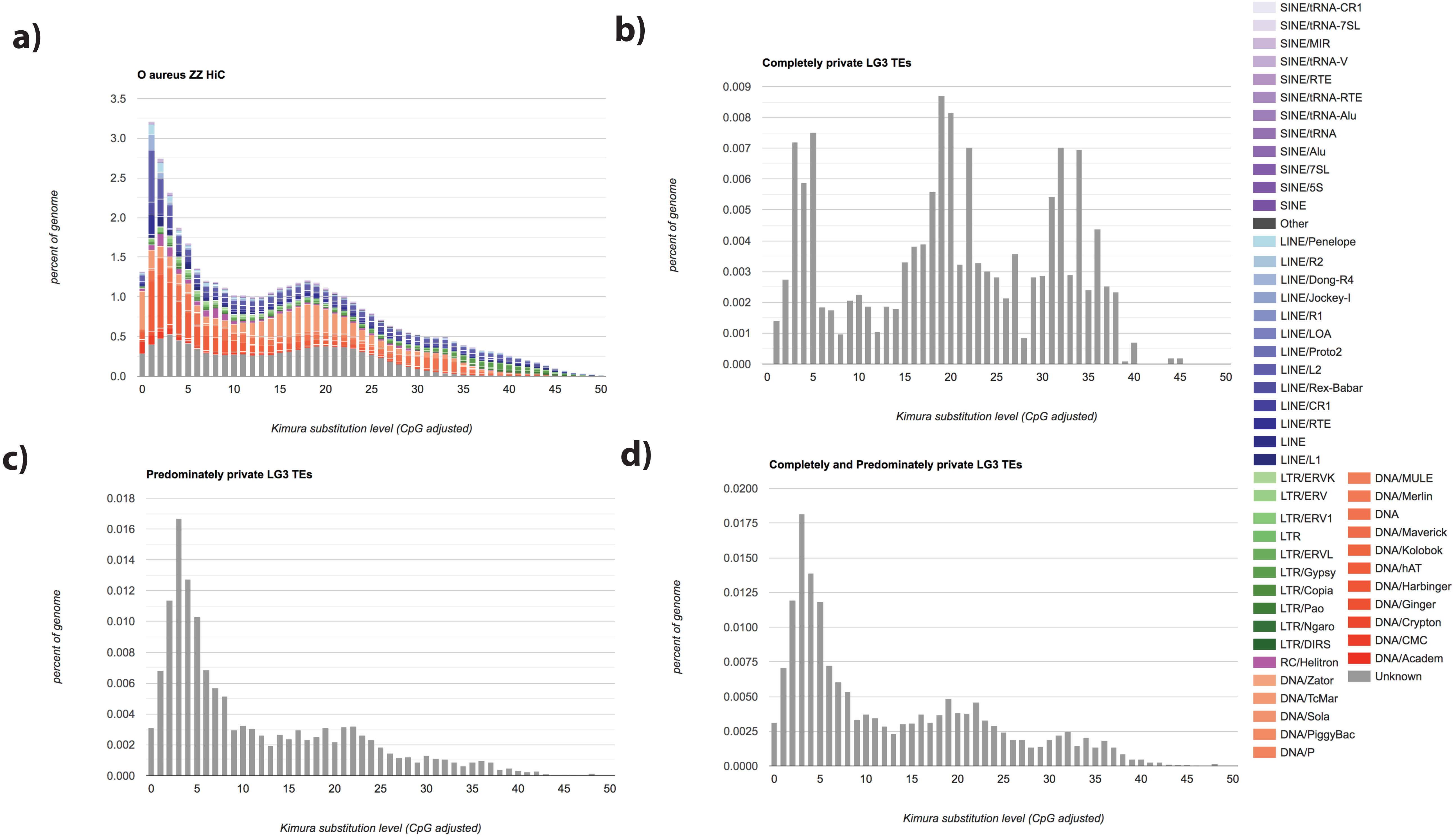
